# ATXN3 regulates lysosome regeneration after damage by targeting K48-K63-branched ubiquitin chains

**DOI:** 10.1101/2025.06.10.656592

**Authors:** Maike Reinders, Bojana Kravic, Pinki Gahlot, Johannes van den Boom, Nina Schulze, Sophie Levantovsky, Stefan Kleine, Markus Kaiser, Yogesh Kulathu, Christian Behrends, Hemmo Meyer

**Affiliations:** Molecular Biology I, Center of Medical Biotechnology, Faculty of Biology, University of Duisburg-Essen; Essen, Germany; Imaging Center Campus Essen, Center of Medical Biotechnology, Faculty of Biology, University of Duisburg-Essen; Essen, Germany; Munich Cluster for Systems Neurology, Medical Faculty, Ludwig-Maximilians-University München; Munich, Germany; Chemical Biology, Center of Medical Biotechnology, Faculty of Biology, University of Duisburg-Essen; Essen, Germany; MRC Protein Phosphorylation and Ubiquitylation Unit, University of Dundee; Dundee, UK

## Abstract

The cellular response to lysosomal damage involves fine-tuned mechanisms of membrane repair, lysosome regeneration and lysophagy, but how these different processes are coordinated is not fully understood. Here we show in human cells that the deubiquitinating enzyme ATXN3 helps restore integrity of the lysosomal system after damage by targeting K48-K63-linked branched ubiquitin chains on regenerating lysosomes. We find that ATXN3 translocates to lysosomes after different types of damage and there colocalizes with its partner, VCP/p97. ATXN3 recruitment occurs late after the initial repair of a subset of lysosomes and after phagophore formation on terminally damaged lysosomes. Of note, inactivation of ATXN3 by induced degradation, depletion or knock-out impairs clearance of damaged lysosomes and full restoration of lysosomal capacity. Mechanistically, ATXN3, along with VCP/p97, turns over K48-K63-branched ubiquitin conjugates on LAMP1-positive, phosphatidylinositol-(4,5)-bisphosphate-decorated compartments that are not yet fully re-acidified indicating involvement in lysosome regeneration. Our findings identify a key role of ATXN3 in restoring lysosomal function after lysosomal membrane damage and uncover K48-K63-branched ubiquitin chain-regulated regeneration as a critical element of the lysosomal damage stress response.

## Introduction

Lysosomal membrane permeabilization (LMP) or full rupture of the limiting membrane of lysosomes and late endosomes constitutes severe cellular stress relevant in various conditions such as neurodegeneration, infection, and cancer. Several conditions cause lysosomal damage including exposure to lysosomotropic compounds, lipid peroxidation and unbalanced lipid compositions associated with neurodegeneration, aging or cancer, as well as cellular uptake of silica, crystals, or pathogens. Cells have developed a complex response, termed the Endo-Lysosomal Damage Response (ELDR), that consists of distinct branches (Meyer & Kravic, 2024; Yang & Tan, 2023; Zoncu & Perera, 2022). Lysosomes with minor damage undergo quick repair of the limiting membrane and re-acidification, which is mediated by the ESCRT machinery and by replenishment of lipids at damage-induced ER-lysosome contact sites (Herbst *et al*, 2020; Radulovic *et al*, 2018; Radulovic *et al*, 2022; Skowyra *et al*, 2018; Tan & Finkel, 2022). If repair fails, individual terminally damaged lysosomes are triaged for destruction by a form of selective macroautophagy, termed lysophagy (Hoyer *et al*, 2022; Hung *et al*, 2013; Maejima *et al*, 2013). In parallel, an mTOR-governed signaling pathway induces the biogenesis of new lysosomal components (Jia *et al*, 2018). In addition, however, a number of recent reports describe, for a subset of lysosomes, damage-induced tubulation and transport events, which are regulated by various factors such as LRRK2, RAB7 or conjugation of ATG8 to single membranes (Bhattacharya *et al*, 2023; Bonet-Ponce *et al*, 2020; Cross *et al*, 2023). Together with the biogenesis of new lysosomal components, they drive the regeneration of functional lysosomes in addition to those that recovered in the initial repair efforts.

A key regulatory feature of the endo-lysosomal damage response is the extensive ubiquitylation of damaged lysosomes (Eapen *et al*, 2021; Kravic *et al*, 2022; Maejima *et al*., 2013) by a number of ubiquitin ligases identified so far (Chauhan *et al*, 2016; Gahlot *et al*, 2024; Liu *et al*, 2020; Teranishi *et al*, 2022; Yoshida *et al*, 2017). An initial wave of ubiquitylation that includes K63-linked ubiquitin chains serves to recruit autophagy receptors and triggers phagophore formation for lysophagy (Eapen *et al*., 2021; Gahlot *et al*., 2024).

However, ubiquitylation regulates additional steps beyond constituting an “eat-me” signal for lysophagy. These functions include the removal of the actin stabilizer CNN2 by the ubiquitin-directed AAA ATPase VCP/p97 to further promote phagophore formation (Kravic *et al*., 2022). Intriguingly, ubiquitylation of lysosomal compartments continues to increase even after phagophore formation around some lysosomes and peaks around 3 h after damage initiation, and this coincides with a further increase in VCP/p97 recruitment (Papadopoulos *et al*, 2017). The role of this late ubiquitylation is as yet unclear, suggesting an unknown function beyond repair and lysophagy in restoring lysosomal functionality.

Using a proteomics approach, we previously identified Ataxin-3 (ATXN3) as a potential regulator in the endo-lysosomal damage response (Koerver *et al*, 2019). ATXN3 is a deubiquitinating enzyme that cooperates with VCP/p97 in various pathways including promoting autophagic flux (Ashkenazi *et al*, 2017; Pfeiffer *et al*, 2017; Wang *et al*, 2006). Mutation of ATXN3 is associated with spinocerebellar ataxia type 3, a hereditary neurodegenerative disease (McLoughlin *et al*, 2020). Here, we show that ATXN3 translocates to lysosomes upon damage and targets conjugates with K48-K63-branched ubiquitin chains on regenerating lysosome, thus revealing a key pathway that helps restore lysosome functionality after damage.

## Results

### ATXN3 translocates to lysosomes in response to lysosomal membrane permeabilization

We first asked whether ATXN3 localizes to lysosomes upon lysosomal damage. To permeabilize lysosomes, we initially applied L-leucyl-L-leucin methyl ester (LLOMe). LLOMe is lysosomotropic and condenses to membranolytic poly-leucine peptides specifically in late endosomes and lysosomes (Thiele & Lipsky, 1990) thus recapitulating damage in pathophysiological conditions. ATXN3-GFP distributed diffusely in unchallenged HeLa cells but robustly translocated to LAMP1-positive compartments upon LLOMe treatment **(Figure 1A,B)**. The translocation occurred relatively late in the damage response with ATXN3 emerging at the end of the 1 h LLOMe treatment and peaking 2 h after LLOMe washout **(Figure 1A,B)**. ATXN3 colocalized with its partner, the AAA-ATPase VCP/p97 **(Figure 1C,D)**, which is an established regulator of the lysosomal damage response (Eapen *et al*., 2021; Klickstein *et al*, 2024; Papadopoulos *et al*., 2017). We confirmed translocation of ATXN3-GFP to damaged lysosomes also in retina pigment epithelium ARPE19 cells **(Figure 1E)**, a non-transformed neuronal cell model (Calcagni *et al*, 2023). ATXN3-GFP was also recruited to lysosomes after treatment with the antihistamine terfenadine **(Supplementary Figure 1A)** which permeabilizes lysosomes (Petersen *et al*, 2013). Moreover, spatially restricted induction of lysosomal membrane permeabilization by light-stimulated lipid peroxidation (Hung *et al*., 2013) led to translocation of ATXN3-GFP to affected lysosomes **(Figure 1F)**.

**Figure 1.**
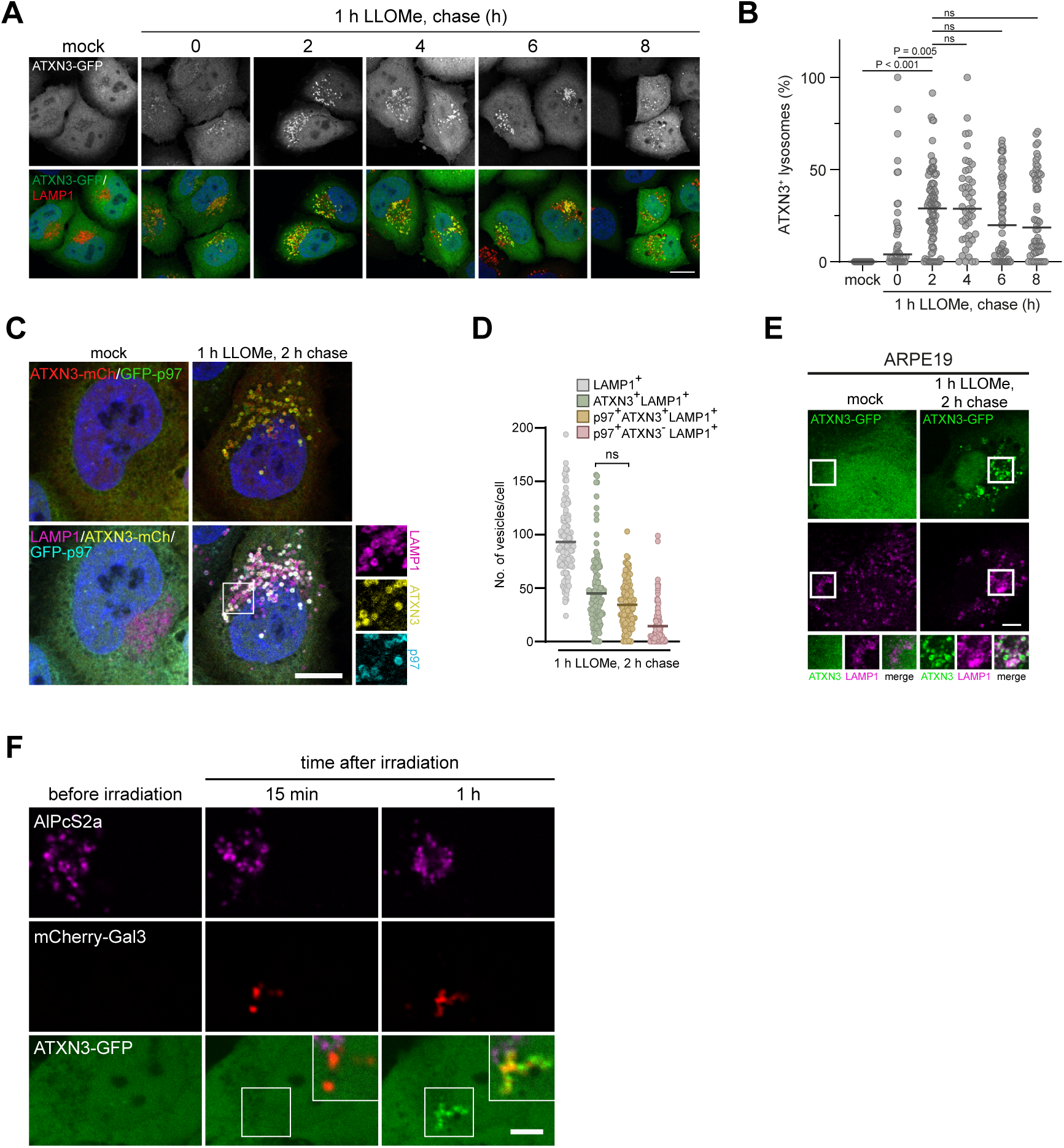
ATXN3 translocates to LAMP1-positive compartments at a late stage after lysosomal damage. (A) HeLa cells expressing ATXN3-GFP were mock or LLOMe-treated (1 mM) for 1 h, fixed at indicated time points after LLOMe washout, and stained with LAMP1 antibodies. Note that ATXN3 translocation peaks at 2 h after washout. Scale bar, 15 µm. (B) Quantification of (A), representative experiment with >40 cells per condition. One-way ANOVA with Dunn’s multiple comparison test. Line indicates median. (C) HeLa cells expressing ATXN3-mCherry and the GFP-p97 (E578Q) substrate-trapping mutant were treated and stained as indicated. Note the robust colocalization on a subset of LAMP1-positive compartments. Scale bar, 10 µm. (D) Quantification of (C). n=3 biological replicates with >30 cells per condition per experiment. One-way ANOVA with Tukey’s multiple comparison test. Line indicates mean. (E) Neuronal ARPE19 cells expressing ATXN3-GFP were mock or LLOMe-treated (1mM) for 1 h followed by 2 h chase before fixation and LAMP1 staining. Scale bar 5 µm. (F) Light-induced lipid peroxidation. Lysosomes in HeLa cells expressing ATXN3-GFP and the damage marker mCherry-Gal3 were loaded with photosensitizer AlPcS2a. Cells were pulse-irradiated and imaged live at indicated time points. Affected lysosomes are bleached and decorated by Gal3. Note recruitment of ATXN3-GFP only after 1h. Scale bar, 5 µm.

Thus, ATXN3 is recruited to lysosomes damaged in various ways to different degrees and in different cell types suggesting that ATXN3 is a general element of the lysosomal damage response.

### ATXN3 is essential for restoring lysosomal degradative capacity after damage

Damaged lysosomes become decorated with cytosolic galectins including galectin-3 (LGALS3, Gal3) that bind to exposed luminal glycans and are easily detectable by immuno-staining (Jia *et al*., 2018; Maejima *et al*., 2013; Thurston *et al*, 2012). The progress of restoring lysosomal integrity during the damage response can be monitored by following the clearance of lysosome-associated Gal3 (Maejima *et al*., 2013). Gal3 clearance occurs initially in those lysosomes that are being repaired and later by lysophagy of a subpopulation of lysosomes that are terminally damaged, as well as by lysosome regeneration (Bhattacharya *et al*., 2023; Eapen *et al*., 2021). We observed a robust dependence of Gal3 clearance on ATXN3, and this was demonstrated by inactivating ATXN3 in three independent ways. We first generated a series of HeLa ATXN3 knockout (KO) cells that showed delayed Gal3 clearance after LLOMe-induced damage **(Supplementary Figure 2A-C)**. The observed delay in Gal3 clearance was rescued by overexpression of wild-type ATXN3, but not of a catalytically inactive ATXN3 C14A mutant demonstrating that ATXN3 function in the lysosomal damage response required its deubiquitinating activity **(Figure 2A,B and Supplementary Figure 2D)**. Likewise, depletion of ATXN3 in HeLa cells by two different siRNAs affected Gal3 clearance **(Figure 2C,D and Supplementary Figure 2E)**. To exclude that the observed effect was indirect due to long-term depletion of ATXN3, we applied a rapid induced-degradation approach. ATXN3 was genomically tagged with FKBP12^F36V^ in U2OS cells **(Supplementary Figure 2F,G)**. Addition of dTAG^VHL^, a proteolysis targeting chimera compound linking FKBP12^F36V^ with the VHL ubiquitin ligase (Nabet *et al*, 2020), induced degradation of ATXN3 efficiently within 1 h **(Supplementary Figure 2H)**. Like in the other approaches, induced degradation of ATXN3 significantly compromised Gal3 clearance after damage infliction **(Figure 2E,F)**. Concurring with results from the other approaches, this demonstrates that ATXN3 is essential for restoring the lysosomal system and its degradative capacity following damage to lysosomal membranes.

**Figure 2.**
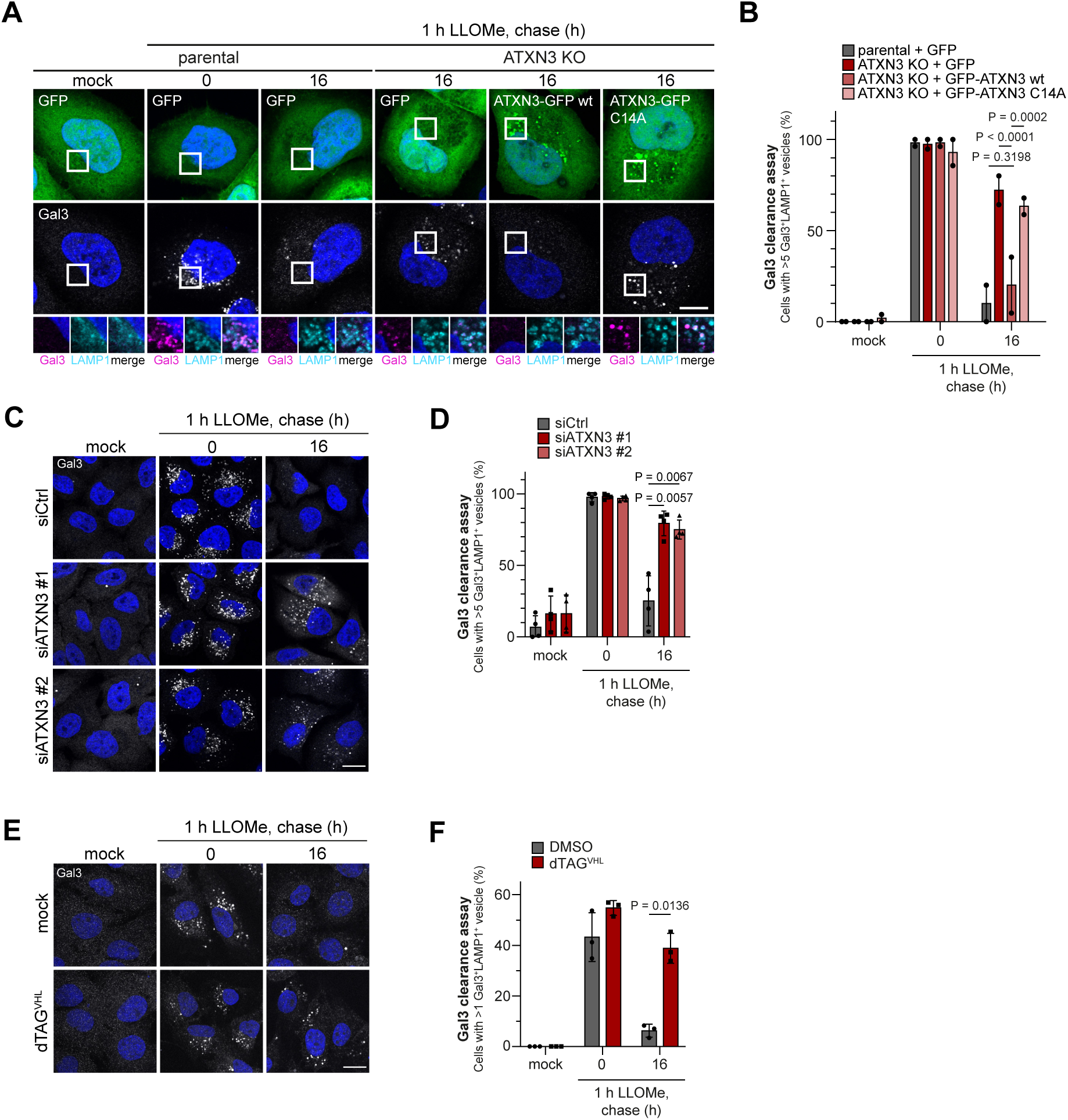
ATXN3 is essential for restoration of degradative compartments after lysosome damage. (A) Knockout-rescue assays for Gal3 clearance. HeLa ATXN3 KO and parental cells expressing indicated constructs were mock or LLOMe-treated (1mM) for 1 h before washout. Cells were fixed at indicated time points and Gal3-positive lysosomes were immuno-stained. Note that Gal3-decorated lysosomes persisted in ATXN3 KO cells, which was rescued by re-expression of ATXN3 wt but not of the catalytically inactive ATXN3 C14A mutant. Scale bar 5 µm. (B) Quantification of (A). n=2 biologically independent experiments with >15 cells quantified per condition per experiment. Error bars, s.e.m. Two-way ANOVA with Tukey’s multiple comparison test was used to test significance. (C) HeLa cells were treated with indicated siRNAs and Gal3 clearance after lysosomal damage assayed as in (A). Scale bar, 15 µm. (D) Quantification of (C). n=4 biologically independent experiments with >30 cells quantified per condition per experiment. Two-way ANOVA with Dunnet’s multiple comparison test was used to test significance. (E) Induced degradation of ATXN3 compromises clearance of damaged lysosomes. The ATXN3 gene was tagged with the FKBP12^F36V^ tag in U2OS cells. ATXN3 degradation was induced by dTAG^VHL^ treatment and Gal3 clearance assessed. Scale bar, 15 µm. See Fig. EV2H for degradation verification. (F) Quantification of (E), n=3 biological replicates with >30 cells per condition per experiment. One-way ANOVA with Šidák’s multiple comparison test.

### ATXN3 is not required for early lysosomal repair pathways

We next aimed to clarify how ATXN3 contributes to lysosomal restoration. Lysosomes with minor damage can be repaired through ESCRT-mediated membrane invagination or by lipid supplementation at ER-lysosome contact sites (Radulovic *et al*., 2018; Radulovic *et al*., 2022; Skowyra *et al*., 2018; Tan & Finkel, 2022). In live cells, we followed initial loss and subsequent recovery of LysoTracker signal which report on the de– and reacidification of damaged lysosomes. Notably, ATXN3 wild-type and KO cells showed the same recovery kinetics **(Figure 3A,B)** indicating that ATXN3 was not required for the early repair pathways that lead to rapid reacidification of a fraction of lysosomes. Consistent with that notion and the late recruitment of ATXN3 observed above, ATXN3-GFP did not largely colocalize with lysosomes positive for the repair factor IST1 as shown by immunostaining of IST1 **(Figure 3C,D)**.

**Figure 3.**
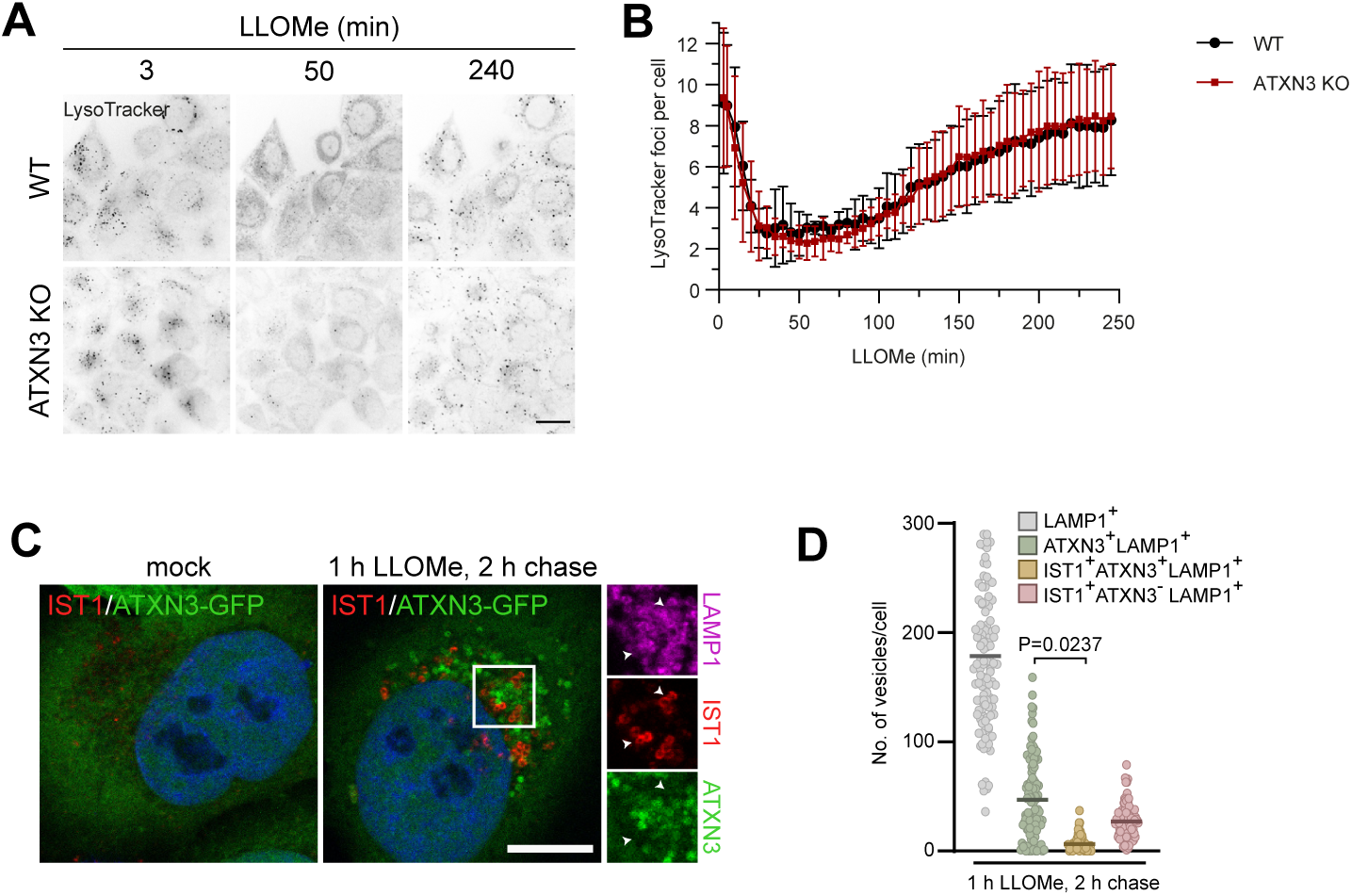
ATXN3 is not involved in early lysosomal repair pathways. (A) HeLa parental and ATXN3 KO cells were loaded with LysoTracker, treated with LLOMe and LysoTracker recovery was imaged over the indicated time period. Note that recovery of LysoTracker is unaffected in ATXN3 KO cells. Scale bar, 20 µm. (B) Quantification of (A), n=3 biologically independent experiments with >70 cells quantified per condition per experiment. (C) Lack of colocalisation of ATXN3 and the ESCRT-III component IST1. HeLa cells expressing ATXN3-GFP were treated as indicated, fixed and immuno-stained for IST1 and LAMP1. Scale bar, 10 µm. (D) Quantification of (C). n=3 biological replicates with >30 cells per condition per experiment. One-way ANOVA with Tukey’s multiple comparison test. Line indicates mean.

### ATXN3 is needed for completion of lysophagy, but not for phagophore formation

We next asked whether ATXN3 was instead involved in lysophagy. To specifically monitor lysophagy, we used a previously established assay that is based on the pH-sensitive mKeima fused to the cytosolic tail of TMEM192 in stable HeLa cell lines (Gahlot *et al*., 2024; Shima *et al*, 2023). Due to its cytosolic location, mKeima reports on acidification in autolysosomes, which enclose the whole lysosome. In contrast, TMEM192-mKeima does not report on acidification of the lysosomal lumen during repair and thus differentiates between the pathways. 6 h after release from LLOMe treatment, control-treated cells showed a shift from green to red-excitable mKeima, indicating a decrease in pH consistent with lysophagy occurring **(Figure 4A and Supplementary Figure 3A)**. In contrast, depletion of ATXN3 largely dampened the increase in red-excitable mKeima, as did depletion of ATG5 or ATG7 as positive controls, indicating a direct function of ATXN3 in lysophagy **(Figure 4A and Supplementary Figure 3A)**. Based on this finding, we speculated that ATXN3 may regulate recruitment of autophagy receptors or phagophore formation. However, depletion of ATXN3 did not affect recruitment of SQSTM1/p62 **(Supplementary Figure 3B,C)** or LC3 **(Figure 4B,C)** to damaged lysosomes compared to control-depleted cells. Moreover, ATXN3-GFP only partially colocalized with LC3 **(Figure 4D,E)**, suggesting that ATXN3 is not involved in phagophore formation but in a later step such as to warrant availability of functional lysosomes for efficient autolysosome formation.

**Figure 4.**
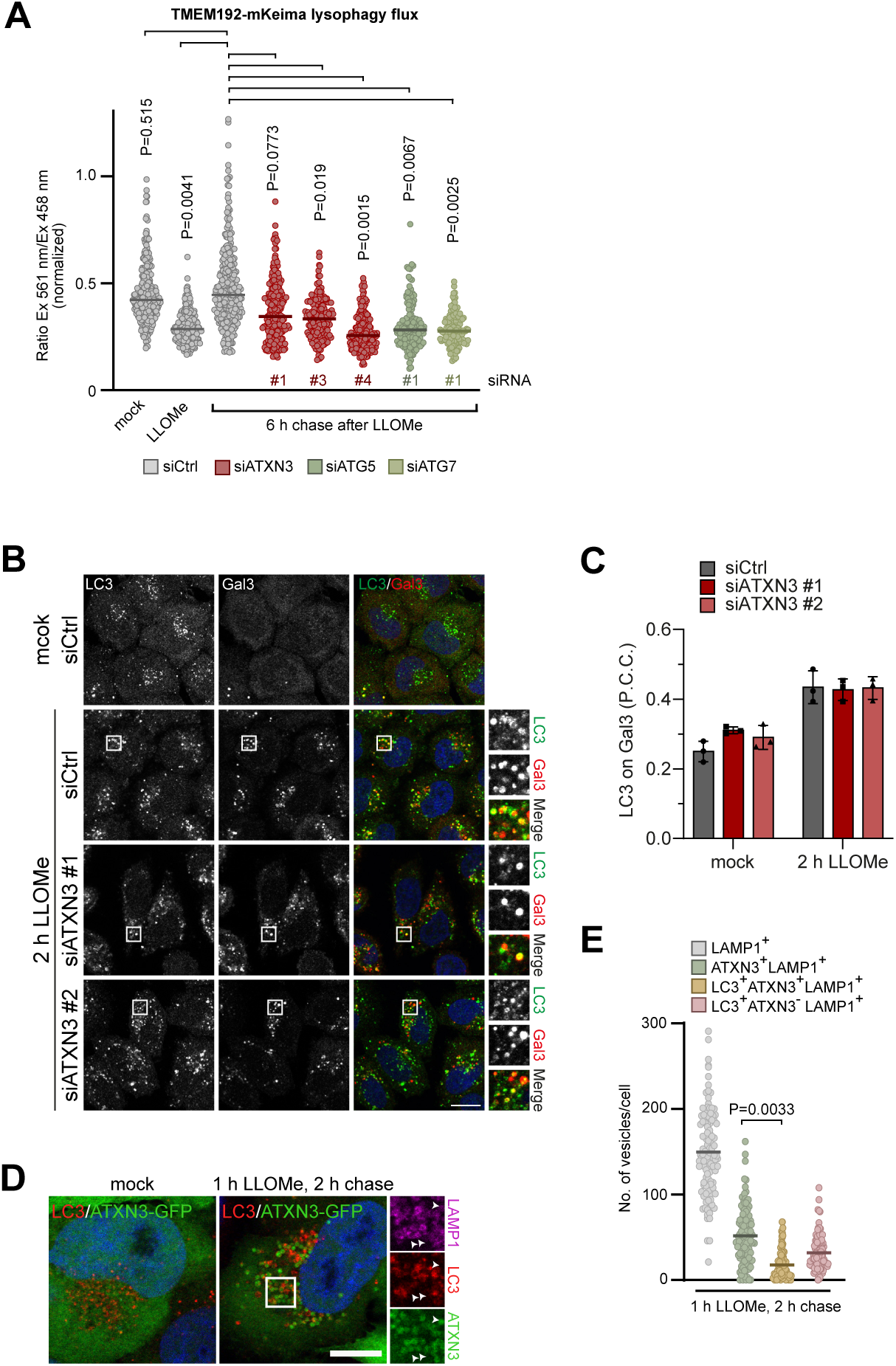
ATXN3 is required for completion of lysophagy, but not for phagophare formation. (A) Lysophagy assay. HeLa cells stably expressing pH-sensitive mKeima fused to the cytosolic terminus of TMEM192. Cells were transfected with indicated siRNAs and treated with LLOMe for 1 h followed by a 6 h washout. mKeima fluorescence was assessed at indicated excitation wavelengths, and the ratio between both intensities determined. Note that the increase in the 561 nm / 458 nm ratio is reduced in ATXN3, ATG5 or ATG7-depleted cells indicating a defect in autolysosome formation. n=4 biological replicates with >30 cells per condition per experiment. One-way ANOVA with Holm-Sidak’s multiple comparison test. Line indicates median. (B) Phagophore formation around damaged lysosomes is not affected by ATXN3 depletion. HeLa cells were treated with indicated siRNAs, incubated with LLOMe before fixation as indicated, and immuno-stained for LC3 and the damage marker Gal3. Scale bar, 15 µm. (C) Quantification of (B). n=3 biologically independent experiments with >70 cells quantified per condition per experiment. (D) Lack of colocalization of ATXN3 with LC3. HeLa cells expressing ATXN3-GFP were mock or LLOMe-treated as indicated, fixed and stained for LC3. Scale bar, 10 µm. (E) Quantification of (D). n=3 biological replicates with >30 cells per condition per experiment. One-way ANOVA with Tukey’s multiple comparison test. Line indicates mean.

### ATXN3 acts on regenerating lysosomes to restore functionality of the lysosomal system

Our results so far indicate that ATXN3 is critical for the restoration of the lysosomal capacity but is not directly involved in the fast initial repair processes of those lysosomes with minor damage or in the formation of phagophores around terminally damaged lysosomes. We therefore explored the localization of ATXN3 in more detail. SIM live-cell microscopy revealed that the majority of ATXN3-decorated LAMP1 compartments were LysoTracker-negative indicating that they were not yet acidified within 3 h since damage infliction **(Figure 5A,B)**. We therefore speculated that ATXN3 localizes to a class of LAMP1 compartments that are subjected to the slower process of regeneration rather than being quickly repaired or autophagocytosed (Bhattacharya *et al*., 2023; Bonet-Ponce *et al*., 2020).

**Figure 5.**
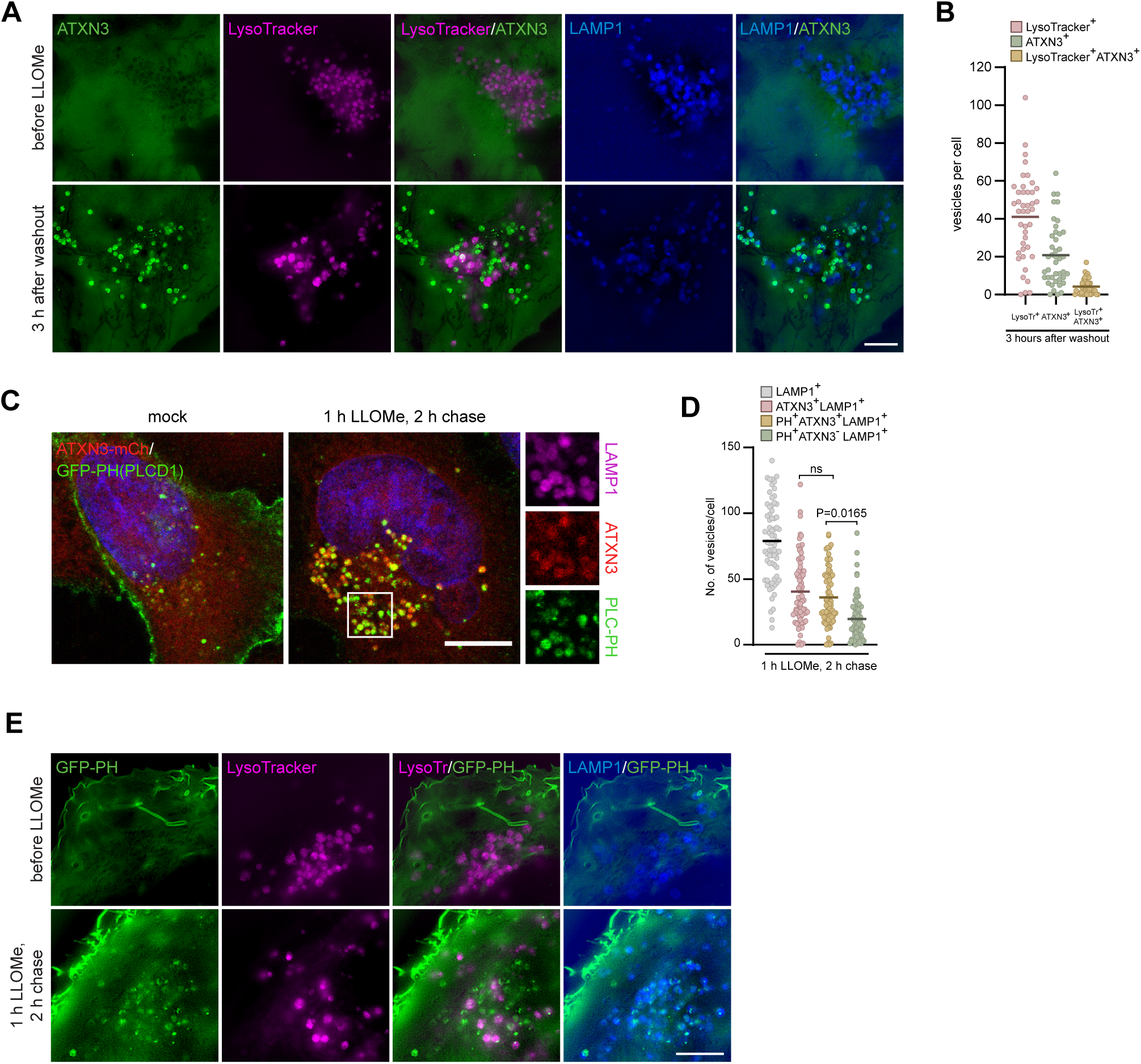
ATXN3 localizes to non-acidified regenerating lysosomes. (A) 3D-SIM live-cell imaging of stable HeLa LAMP1-BFP cells loaded with lysotracker and expressing ATXN3-GFP. Cells were treated with 1mM LLOMe for 12 minutes, washed and followed during recovery in medium containing lysotracker for 3 h. Scale bar 5 µm. (B) Quantification of 2D-SIM images taken in experiments shown in (A). n=2 biologically independent experiments with >20 cells quantified per experiment. Lines indicate median. (C) HeLa cells expressing ATXN3-mCherry and the PI(4,5)P2 sensor GFP-PH(PLCD1) were mock or LLOMe-treated as indicated and immuno-stained for LAMP1. Note the colocalization of ATXN3 and GFP-PH on a subpopulation of lysosomes. Scale bar, 10 µm. (D) Quantification of (C), n=3 biological replicates with >20 cells per condition per experiment. One-way ANOVA with Tukey’s multiple comparison test. Line indicates mean. (E) SIM live-cell imaging of HeLa cells expressing LAMP1-BFP and indicated reporters and treated with 1 mM LLOMe as indicated. Scale bar 5 µm.

Phosphatidylinositol (4,5)-bisphosphate (PI(4,5)P_2_) has been implied as a marker for regenerating lysosomes based on recruitment on clathrin-related proteins (Bhattacharya *et al*., 2023). We used the pleckstrin homology (PH) domain of PLCD1 fused to GFP as an established PI(4,5)P_2_ sensor, which localized to the plasma membrane in control cells, as expected (Stauffer *et al*, 1998). Of note, the PH-GFP sensor translocated to ATXN3 and LAMP1-positive compartments specifically after lysosomal damage induction **(Figure 5C,D)**. Live-cell microscopy with LysoTracker confirmed that these LAMP1-positive compartments decorated with PH-GFP were largely not re-acidified even 3 h after damage infliction **(Figure 5E)**. Thus, ATXN3 primarily acts on LAMP1-positive, yet unacidified compartments, which are regenerating rather than being subjected to lysophagy, and thereby helps restore the functionality of the lysosomal system. This is consistent with the delay in Gal3 clearance observed above in ATXN3-deficient cells and is in line with a reduced recovery of lysosomal degradative capacity after damage in ATXN3 KO cells as monitored by the fluorogenic proteolysis reporter DQ-BSA in lysosomes **(Supplementary Figure 4A)**.

### ATXN3 and VCP/p97 target damage-induced K48-K63 branched ubiquitin conjugates on regenerating lysosomes

We next aimed to address the molecular basis for ATXN3 function in the lysosomal damage response. ATXN3 was recently found to specifically bind and cleave ubiquitin chains with branched K48 and K63 linkages and to cooperate with VCP/p97 to process these conjugates (Lange *et al*, 2024). We used a nanobody, NbSL3.3Q that specifically detects K48-K63-branched ubiquitin chains (Lange *et al*., 2024) to ask whether K48-K63 chains played any role in ELDR. Of note, over-expressed NbSL3.3Q-GFP colocalized with lysosomes specifically after lysosomal damage indicating that K48-K63-branched ubiquitin chains are a signal in the lysosomal damage response **(Supplementary Figure 5A,B)**.

Given the high affinity of NbSL3.3Q for branched chains, we wanted to avoid interfering with the dynamics of K48-K63-branched ubiquitin chains due to NbSL3.3Q binding in live cells, and therefore used chromophore-conjugated NbSL3.3Q for immunostaining of fixed cells.

The staining revealed that K48-K63 chains peaked late during the damage response at 2 h chase and decreased over the full 8 h of analysis **(Figure 6A,B)** comparable to the ATXN3 dynamics shown above. Importantly, co-imaging showed that K48-K63-branched chains and ATXN3 largely colocalized on LAMP1-positive compartments **(Figure 6C,D)**. Of note, the K48-K63-branched chains also colocalized with p97 **(Figure 6E,F)** suggesting that p97 and ATXN3 cooperate in targeting K48-K63 conjugates. Indeed, chemical inhibition of p97 or knockout of ATXN3 led to an enhanced accumulation and persistence of K48-K63-branched chains as visualized by NbSL3.3Q staining showing that ATXN3 in cooperation with p97 turns over the K48-K63-branched ubiquitin conjugates to regulate restoration of lysosomal capacity **(Figure 6G-J)**. This indicates that processing of K48-K63-branched ubiquitin chains is essential for cells to cope with the stress induced by lysosomal damage. Thus, the conjugation of LAMP1-positive compartments with K48-K63-linked ubiquitin chains and the subsequent turnover by ATXN3 and p97 is an integral part of the clearance and regeneration processes that restore lysosomal capacity after damage.

**Figure 6.**
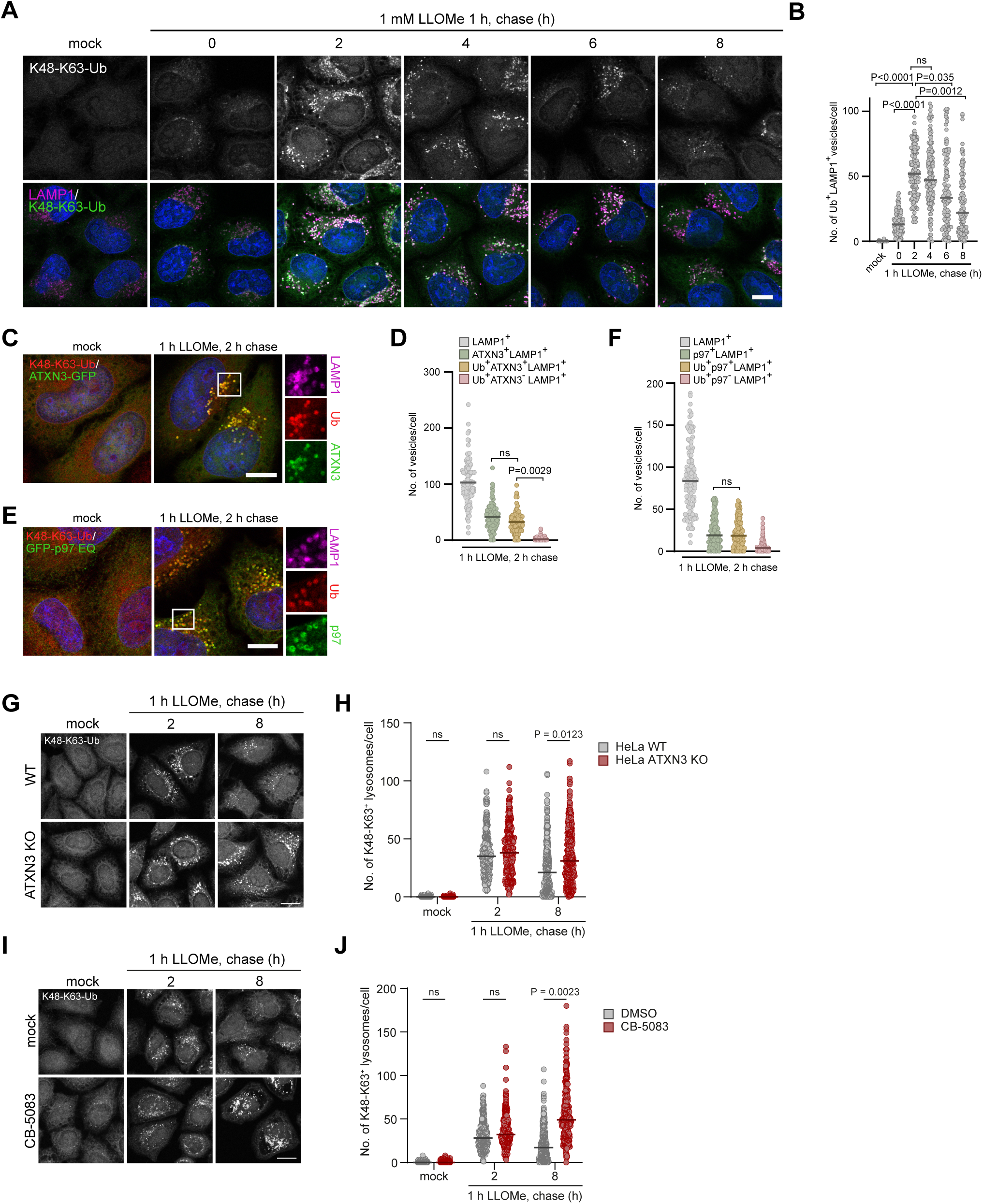
ATXN3 and VCP/p97 target and turn over damage-induced K48-K63 branched ubiquitin conjugates on regenerating lysosomes. (A) HeLa cells were LLOMe-treated and chased for indicated periods of time. Cells were fixed and immuno-stained with LAMP1 antibodies and an AlexaFluor568-conjugated nanobody NbSL3.3Q specific for K48-K63-branched ubiquitin chains. Note the nanobody signal peaking at 2 h chase on a subpopulation of lysosomes and fading off within 8 h. Scale bar, 10 µm. (B) Quantification of (A), n=3 biological replicates with >30 cells per condition per experiment. One-way ANOVA with Holm-Sidak’s multiple comparison test. Line indicates mean. (**C and E**) HeLa cells expressing ATXN3-GFP (C) or p97-E578Q (E) were fixed and stained for K48-K63-branched ubiquitin chains with NbSL3.3Q. Scale bar, 10 µm. (**D and F**) Quantification of (C and E), n=3 biological replicates with >30 cells per condition per experiment. One-way ANOVA with Tukey’s multiple comparison test. Line indicates mean. (G) Parental or ATXN3 KO cells were pulse-treated with LLOMe, chased for indicated times and stained with NbSL3.3Q. Note persistence of K48-K63-branched chains in ATXN3 KO cells. Scale bar, 15 µm. (H) Quantification of (G) n=3 biological replicates with >50 cells per condition per experiment. Two-way ANOVA with Šidák’s multiple comparison test. Line indicates median. (I) HeLa cells were incubated with p97 inhibitor CB-5083 or vehicle alone and LLOMe treated and processed as in (G). Scale bar, 15 µm. (J) Quantification of (I) n=3 biological replicates with >70 cells per condition per experiment. Two-way ANOVA with Šidák’s multiple comparison test. Line indicates median.

## Discussion

In this study, we uncover a key role of ATXN3 in the cellular response to lysosomal membrane permeabilization. By analyzing cancer and neuronal cell models, we find that compromising ATXN3 function leads to a severe delay in clearing and regenerating damaged lysosomes and thus in restoring functionality of the lysosomal system after lysosomal damage. The fact that ATXN3 translocates to lysosomes in response to damage and that it processes stress-induced ubiquitin-conjugates on lysosomal membranes indicates that the function of ATXN3 is direct. Previous work showed that ATXN3 regulates the stability of the PI3 kinase class III complex component beclin-1/BECN1, which triggers phagophore formation, and that ATXN3 thus supports long-term autophagic flux (Ashkenazi *et al*., 2017; Hill *et al*, 2021). This function does not appear to be directly relevant in our setup that monitors acute stress of lysosomal damage, because we find that LC3 lipidation and LC3 decoration of damaged lysosomes during lysophagy is not severely affected in ATXN3-compromised cells. Thus, ATXN3 has an additional critical function in ensuring cellular homeostasis by restoring integrity of the lysosomal system in stress conditions.

Mechanistically, we demonstrate that ATXN3 fulfils its function by targeting and processing ubiquitin conjugates with K48-K63-branched ubiquitin chains. K48-K63-branched chains have been established as a stress response signal only recently (Lange *et al*., 2024). Our work shown here now confirms their relevance in stress signaling and puts K48-K63-branched chains at the center of the lysosomal damage response. Notably, we show that ATXN3 cooperates with its partner, the ubiquitin-directed AAA ATPase VCP/p97, in processing K48-K63-branched ubiquitin conjugates. VCP/p97 is an important regulator of many ubiquitin-regulated stress response pathways in which VCP/p97 mobilizes and unfolds ubiquitylated client proteins, often for subsequent degradation in the proteasome (van den Boom & Meyer, 2018; Ye *et al*, 2017). The established targets of VCP/p97 are K48-linked conjugates (Bodnar & Rapoport, 2017; Olszewski *et al*, 2019). It is possible that, during the damage response, K48-linked conjugates are being modified with additional K63-linked branches in order to prioritize them over the bulk of other VCP/p97 targets. The role of ATXN3 could be to preferably direct VCP/p97 to these targets and then apply the biochemically established debranching activity of ATXN3 to render them palatable for subsequent mobilization and unfolding by VCP/p97.

On the cell biological level, our work, with the identification of a pathway regulated by ATXN3 and K48-K63-branched ubiquitin chains, puts a spotlight on regenerating lysosomes.

Research has largely focused on the early repair pathways driven by ESCRT machinery and lipid replenishment, as well as on lysophagy of terminally damaged lysosomes (Bohannon & Hanson, 2020; Hoyer *et al*., 2022; Yang & Tan, 2023; Zhen *et al*, 2021). Recent work has identified additional mechanisms that are critical for the response to lysosomal damage but that do not directly mediate membrane resealing or lysophagy, suggesting functions in lysosome regeneration. One is the role of a tubulation process that resembles autophagic lysosome regeneration (ALR) in regenerating damaged lysosomes (Yu *et al*, 2010) (Bhattacharya *et al*., 2023). Moreover, the Parkinson’s disease-associated kinase LRRK2 was found to regulate tubulation of damaged lysosomes (Bonet-Ponce *et al*., 2020). So far, how these pathways contribute to regeneration of lysosomes after damage infliction is not fully understood. One possibility is that regeneration could retrieve lysosomal membranes from damaged lysosomes for the biogenesis of new lysosomes. Conversely, the regeneration process could remove material that is damaged or not needed from lysosomes before these lysosomes re-acidify and resume their functionality. In a not mutually exclusive scenario, it is also possible that the regenerating lysosomes are acceptor compartments for new material from the biosynthetic pathway or for recycled material from canonical ALR. Importantly, our data highlight that regenerating lysosomes represent a considerable fraction of LAMP1-positive compartments during the damage response and that the underlying processes are highly regulated in unanticipated ways involving K48-K63-branched ubiquitin chains and their processing by ATXN3 and p97. With ATXN3 and p97 being linked to neurodegeneration (McLoughlin *et al*., 2020; Meyer & Weihl, 2014), our findings underscore the relevance of this part of the lysosomal damage response for maintaining cellular homeostasis.

## Author contributions

**Maike Reinders, Bojana Kravic, Pinki Gahlot:** Conceptualization; Formal analysis; Investigation; Methodology; Data curation; Visualization; **Johannes van den Boom:** Investigation; **Nina Schulze:** Data curation; Methodology; Investigation; Funding acquisition; **Sophie Levantovsky:** Investigation; **Stefan Kleine:** Resources; **Markus Kaiser:** Conceptualization; Resources; Funding acquisition; **Yogesh Kulathu:** Conceptualization; Resources; Funding acquisition; **Christian Behrends:** Conceptualization; Resources; Funding acquisition; **Hemmo Meyer:** Conceptualization; Data curation; Formal analysis; Supervision; Funding acquisition; Visualization; Writing—original draft; Project administration; Writing—review and editing.

## Acknowledgments

We acknowledge the use of equipment and the support by the Imaging Center Campus Essen (ICCE), and specifically thank J. Koch for his help. This research was funded by a joined grant of the Deutsche Forschungsgesellschaft (DFG, German research foundation) to H.M. and C.B. (Project-ID 447112704). H.M., M.K. and N.S. were supported by the DFG CRC 1430 (ID 424228829), and C.B. by EXC 2145 SyNergy (ID 390857198) and the CRC 1177 (ID 259130777). Microscopes were funded by DFG INST 20876/480-1 FUGG and INST 20876/294-1 FUGG. Y.K. was funded by ERC Consolidator grant (StressHUb grant 101002428) and MRC grant MC_UU_00038/3.

## Methods

### Plasmids

pEGFP-N1-ATXN3 and pmCherry-N1-ATXN3 plasmids were created from the plasmid pcDNA5FRT/TO-Ataxin3-Strep-HA (Hulsmann *et al*, 2018) using Gibson cloning. To generate the catalytically inactive form of ATXN3, site-directed mutagenesis was performed to obtain pEGFP-N1-ATXN3-C14A. Homology arms for the N-terminus of ATXN3 (with silent mutations to prevent sgRNA binding) were ordered as a gBlock (Integrated DNA Technologies) containing HindIII and BamHI restriction sites. The gBlock was cloned into pBlueScript-SK(+) vector by restriction digestion. PuroR-P2A-HA-dTAG from pCRIS-PITChv2-Puro-dTAG-BRD4 plasmid (gift from James Bradner; addgene #91793) was inserted into the homology arms by Gibson cloning. Nanobody expression plasmids were described before (Lange *et al*., 2024). GFP-C1-PLCdelta-PH plasmid was a gift from Tobias Meyer (addgene #21179). pEGFP-C1-p97 E578Q plasmid was described before (Kravic et al., 2022). LAMP1 was cloned into pcDNA3-eBFP2 vector by amplifying human LAMP1 from HeLa cDNA using oligos containing HindIII and BamHI sites.

### Cell lines

HeLa ATXN3 knockout cells were generated by transfection with Ataxin-3 Double Nickase Plasmid (SantaCruz; sc-417498-NIC) according to manufacturer’s instructions. U2OS ATXN3^dTAG^ cells were generated using sgRNA (ACTCACTTTCTCGTGGAAGA) targeting the N-terminus of ATXN3 using CRIPSR Cas9 technology to allow integration of PuroR-P2A-HA-dTAG (derived from pCRIS-PITChv2-Puro-dTAG-BRD4, Addgene #91793). Cells were cultured in DMEM containing 10% fetal bovine serum (FBS, PAN-Biotech), and 1% Penicillin/Streptomycin (PAN-Biotech) and selection antibiotic Puromycin (1 µg/ml).

Integration was confirmed by Sanger sequencing following genomic PCR. To amplify the N-terminus of ATXN3 from genomic DNA in order to confirm the integration the following primers were used: 5’-CCCCGTCTCCCACACAATTTA-3’ and 5’-CAGCAGGCTAGGCAGACTAC-3’. The primers 5’-TTCACTCGCTCTTCGCTTCA-3’, 5’-AGTTCTTGCAGCTCGGTGAC-3’; 5’-ATCATCCCACCACATGCCAC-3’, 5’-AAGCGATGGAAAGTGACGGA-3’ were used for Sanger sequencing and integration of PuroR-P2A-HA-dTAG at the N-terminus of ATXN3 was confirmed. The knock-in was also confirmed via western blot. For generation of HeLa LAMP1-BFP stable cell line, HeLa Kyoto cells were transfected with circular pcDNA3-Puro-LAMP1-eBFP2 and Puromycin (1 µg/ml).

### Cell culture

HeLa and U2OS cells were cultured in DMEM containing 10% FBS and 1% Penicillin/Streptomycin. ARPE19 cells were cultured in DMEM:F12 supplemented with 10% FBS and 1% Penicillin/Streptomycin. Cells were grown in standard conditions, 37 °C and 5% CO_2_. Cells were transfected with plasmids using Lipofectamine 2000 (ThermoFisher Scientific) or with 10 nM siRNA using RNAiMAx (ThermoFisher Scientific) according to the manufacturer’s instructions. Transfected cells were analyzed after 24 h (plasmids) or 48 h (siRNA).

### RNA interference

siRNAs targeting ATXN3 were purchased from Microsynth: siATXN3 #1

UGGCAGAAGGAGGAGUUACTT (Wang *et al*, 2012), siATXN3 #2

CAGGGCUAUUCAGCUAAGUAUTT (Sacco *et al*, 2014), siATXN3 #3

GCACUAAGUCGCCAAGAAATT (Ashkenazi *et al*., 2017), siATXN3 #4 GCAGGGCUAUUCAGCUAAGTT (Ashkenazi *et al*., 2017).

### Cell treatments

To induce lysosomal damage, cells were treated with 1 mM L-Leucyl-L-leucine methyl ester hydrobromide (LLOMe, Sigma) or in the case of HeLa TMEM192-mKeima with 500 µM LLOMe for indicated times or with 8 µM Terfenadine for 24 h. For p97 inhibition, cells were treated with 5 µM CB-5083 (Selleckchem, #S8101) for indicated times. To induce degradation of ATXN3^dTAG^, cells were treated with 1 µM dTAG^VHL^ (Torcis, #6914) for indicated times.

### Antibodies

The following primary antibodies were used in the study: anti-Ataxin-3 (mouse, BioLegend, 650402; WB 1:1000), anti-Galectin-3 (rat, Santa Cruz Biotechnology, sc-23938; IF 1:500), anti-LAMP1 (mouse, Santa Cruz Biotechnology, sc-20011; IF 1:500), anti-LAMP1 (rabbit, Cell Signaling, D2D11; IF 1:500), anti-LC3 (rabbit, MBL, PM036; IF 1:500), anit-p62/SQSTM1 (mouse, Abonva, H00008878-M01; IF 1:500), anti-GFP (mouse, Santa Cruz Biotechnology, sc-9996; IF 1:500), anti-GFP (mouse, Roche, 11814460001; IF 1:500), anti-IST1 (mouse, Proteintech, #66989, IF 1:500), anti-Tubulin (mouse, Sigma, T-5168; WB 1:8000. HRP-coupled secondary antibodies were purchased from Bio-Rad and Alexa Fluor-conjugated secondary antibodies from Invitrogen.

### Immunofluorescence staining and confocal laser-scanning microscopy

NbSL3.3Q nanobody was expressed in *E. coli* BL21 and purified as described (Lange et al., 2024). The purified nanobody was labelled with Alexa Fluor 568 NHS ester (Lumiprobe; 8-fold molar excess) and repurified using a Superdex 75 10/300 column (GE Healthcare).

Cells were cultured on coverslips, fixed in 4% paraformaldehyde at room temperature or in 100% methanol at –20 °C (for NbSL3.3Q staining), permeabilized with 0.1% Triton X-100 in PBS and blocked in PBS with 3% BSA + 0.1% Triton X 100 + 0.1% saponin. Following immunofluorescence staining, cells were mounted in ProLong Gold (Thermo Fisher Scientific).

Confocal laser scanning microscopy was performed on a Leica TCS SP8X Falcon confocal microscope (Leica Microsystems), equipped with HyDs SMD detectors, HC PL APO 63x/1.4 Oil CS2 objective, white-light laser, Argon laser and a 405 diode laser and the Leica Application Suite X (LAS-X) software version 3.5.7. Images were acquired in a 1024×1024 ox format (1.5x zoom) with a bit depth of 12 bit.

In additional, imaging was performed on a TCS SP8 HCS A confocal microscope using the LAS X software (3.5.7.23225) An HC PL APO 63×/1.4 NA CS2 oil immersion objective was used to acquire the images. Fluorophores were excited with a 405 nm diode laser, an argon laser line 488 nm, a 561 nm diode-pumped solid-state laser, and a 633 nm helium-neon laser. Signals were detected with standard PMT detectors and sensitive Hybrid detectors (HyD) in a 2048×2048 px format (1x zoom) with a bit depth of 12 bit.

### Live cell microscopy, photodamage and SIM

Live cell imaging was performed at 37 °C in imaging medium (P04-03591, PAN-Biotech) supplemented with 10% FCS and L-Glutamine (11500626, Fisher Scientific). For lysotracker fluorescence intensity experiment, cells were incubated with LysoTracker Deep Red (Thermo Fisher) for 1 h and washed twice with PBS before addition of imaging medium.

Prior to treatment with LLOMe, multiple regions were defined per sample for automated imaging using an Eclipse Ti-E (Nikon) inverted microscope equipped with an Andor AOTF Laser Combiner, a CSU-X1 Yokogawa spinning disk unit and an iXon3 897 single photon detection EMCCD camera (Andor Technology). CSU 640 nm laser and CFI Apo TIRF 60x/1.49 oil immersion objective (Nikon) were used for image acquisition. Every region in each sample was imaged once before addition of LLOMe and then every 5 minutes after LLOMe treatment to track loss of lysotracker fluorescence intensity from lysosomes.

Spinning disk microscopy with laser induced lysosomal damage was performed on the Eclipse Ti-E (Nikon) inverted microscope with an Andor ILE Laser Combiner, a CSU-X1 Yokogawa spinning disk unit and a ZL41 Cell sCMOS camera (Andor Technology). Laser lines used for excitation of EGFP and mCherry were 488 nm and 561 nm, respectively.

Images were acquired using a CFI APO TIRF 60x/1.49NA oil immersion objectives (Nikon). To induce LMP with light, cells were treated with 125 nM AlPcS2a (P40632, Frontier Scientific) as described before (Kravic et al., 2022). Damage was induced with a 640 nm laser using Micropoint 4 Scanned Photo Stimulation device (Andor Technology). Image acquisition was controlled by Andor IQ3 Software (Andor Technology) as described before (Kravic et al., 2022).

HeLa TMEM192-mKeima cells were plated in 8-well Ibidi µ-slide and transfected with indicated siRNAs. After 48 h of knockdown, live cells were imaged before and after 1 h of treatment with 500 µM LLOMe using Leica TCS SP8X Falcon confocal microscope (specifications above) at 37 °C in imaging medium. Cells were washed twice with PBS after LLOMe treatment, maintained in imaging medium for recovery and imaged again after 6 h of chase.

Images were processed using Fiji software (https://imagej.net/Fiji), Adobe Photoshop, and Illustrator. Automated image analysis was done with CellProfiler software version 2.1 (Carpenter *et al*, 2006; Stirling *et al*, 2021). Excel 2016 (Microsoft Corporation) and GraphPad Prism 10.3 (GraphPad Software Inc.) were used for graphical representation and statistical analysis.

### Structured Illumination Microscopy (SIM)

Live HeLa LAMP1-BFP cells were imaged at 37 °C and 5% CO_2_ in imaging medium (P04-03591, PAN-Biotech) supplemented with 10% FBS and 1% Penicillin/Streptomycin. Cells were transfected with plasmid encoding ATXN3-GFP or GFP-PLC PH one day before imaging. The cells were loaded with LysoTracker Deep Red (LysoTr) for 10 min prior to imaging. Live cells were imaged once before LLOMe treatment, then treated with 1 mM LLOMe for 12 min or 1 h, washed twice with PBS and incubated in imaging medium containing LysoTracker to track recovery of LysoTracker and recruitment of the overexpressed protein. Cells were imaged again after washout at indicated time points.

For live cell imaging using 2D-SIM and 3D-SIM, images were acquired using a commercial Nikon N-SIM S microscope system equipped with ORCA-Fusion BT sCMOS camera (Hamamatsu Photonics K.K.) using NIS Elements software (Nikon). Images were acquired using a CFI SR HP Apochromat TIRF 100xAC (NA 1.49) oil immersion objective. ZIVA light engine (Lumencor) equipped with six solid-state laser lines was used as excitation light source. Laser lines used for excitation of Lysotracker Deep Red, GFP and BFP were 637 nm, 476 nm and 405 nm, respectively. Single-band bandpass emission filters FF01-460/60 for BFP, FF01-525/50 for GFP and FF01-692/40 for Lysotracker Deep Red (Semrock) were used. Multipoint timelapse images were acquired in three individual z-planes per position with z-spacing distance of 0.5 µm. 2D-SIM and 3D-SIM raw data were reconstructed using the Slice Reconstruction in NIS-Elements with default settings.

### Lysotracker washout Analysis 2D-SIM

Automated image analysis of 2D-SIM LLOMe washout experiments was performed with CellProfiler software (version 4.2.6) (Stirling *et al*., 2021). Cells were identified using the Run Omnipose plugin for CellProfiler with the cyto2 model (Cutler *et al*, 2022). Accuracy of cell detection and, if necessary, correction of cell outlines was controlled by a manual editing step (EditObjectsManually). To enhance vesicular structures of ATXN3, LAMP1and LysoTracker, feature enhancement using Difference of Gaussians (DoG) was applied to the respective channels. Vesicles were detected and related to the respective parent cells. Double-positive vesicles were detected using the MaskObjects module with a minimum overlap fraction of 0.2.

### DQ-BSA Assay

Cells were treated with DQ-BSA reagent (DQ™ Red BSA (Invitrogen™) Cat.No D12051) for 7 h at concentration of 10 µg/ml. Prior to microscopy, cells were fixed in PFA, stained with DAPI and mounted in ProLong Gold (Thermo Fischer Scientific). Samples were imaged using a Leica TCS SP8X Falcon confocal microscope (Leica Microsystems) as described above.

### Immunoblotting

Cells were lysed lysis buffer consisting of 150 mM KCl, 50 mM Tris-HCl pH 7.4, 5 mM MgCl_2_, 5% Glycerol, 1% Triton X-100, 2 mM ß-Mercaptoethanol and supplemented with protease inhibitors (complete EDTA-free protease inhibitor cocktail, Roche). Proteins were separated by SDS–PAGE and transferred to nitrocellulose membranes (Amersham, GE Healthcare). Immunoblot analysis was performed with the indicated antibodies and visualized with SuperSignal West Pico Chemiluminescent substrate (Pierce).

### Statistical analysis

All quantitative data are presented as the mean ± s.d. of biologically independent samples or all data points from independent experiments are presented with the line indicating median, unless stated otherwise. Statistical analysis was carried out using GraphPad Prism 10.3 software on the mean of independent experiments. One– or two-way ANOVA was used to determine statistical significance, unless stated otherwise. A p-value <0.05 was considered statistically significant.

