## Supplemental figures and legends for "ATXN3 regulates lysosome regeneration after damage by targeting K48-K63-branched ubiquitin chains"

**Supplementary figures**

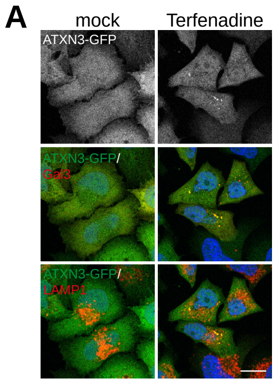


Supplementary Figure 1

**Supplementary Figure 1 (related to Fig. 1). ATXN3 is recruited to lysosomes damaged by terfenadine.**

(**A**) HeLa cells expressing ATXN3-GFP were treated with 8 μM terfenadine for 24 h, fixed and immuno-stained for Gal3 and LAMP1. Scale bar, 15 μm.

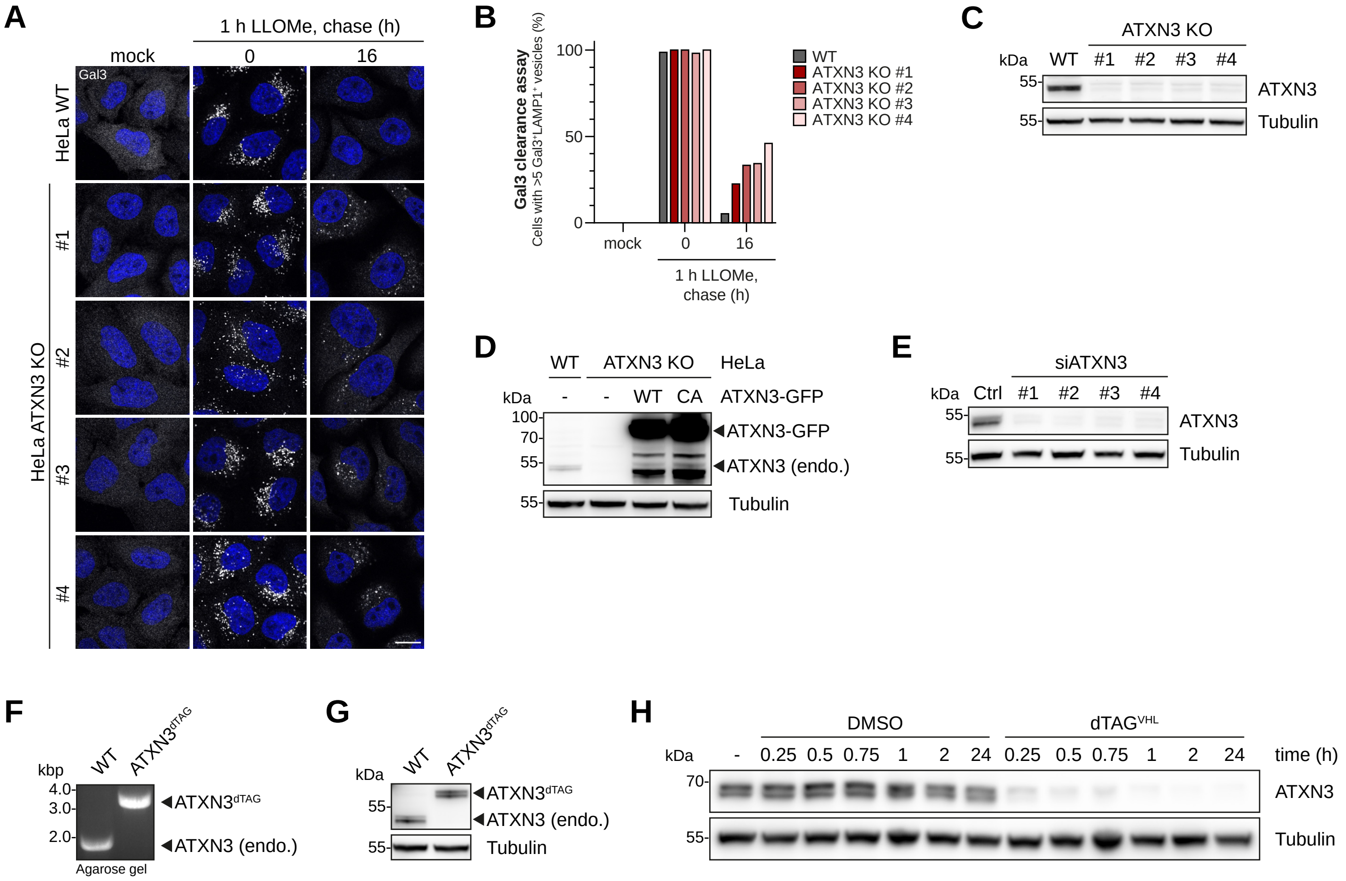


Supplementary Figure 2

**Supplementary Figure 2 (related to Fig. 2): ATXN3 is essential for clearance of damaged lysosomes and restoration of degradative compartments after lysosome damage.**

(**A**) Gal3 clearance assays in HeLa parental and 4 different ATXN3 KO clones performed as in Fig. 2A. Scale bar, 15 µm.
(**B**) Quantification of (A), representative experiment with >40 cells per conditions.
(**C**) Western blot verification of ATXN3 KO.
(**D**) Western blot verification of rescue constructs shown in Fig. 2A.
(**E**) Western blot evaluation of siRNA-mediated ATXN3 depletion.
(**F**) PCR on genomic DNA confirming genomic insertion of the $FKBP^{F36V}$ tag in $ATXN3^{dTAG}$ U2OS cells.
(**G**) Western blot analysis of lysates of parental U2OS (WT) and gene-edited $ATXN3^{dTAG}$ cells.
(**H**) Western blot assessment of the induced degradation of $ATXN3^{dTAG}$ in U2OS cells.

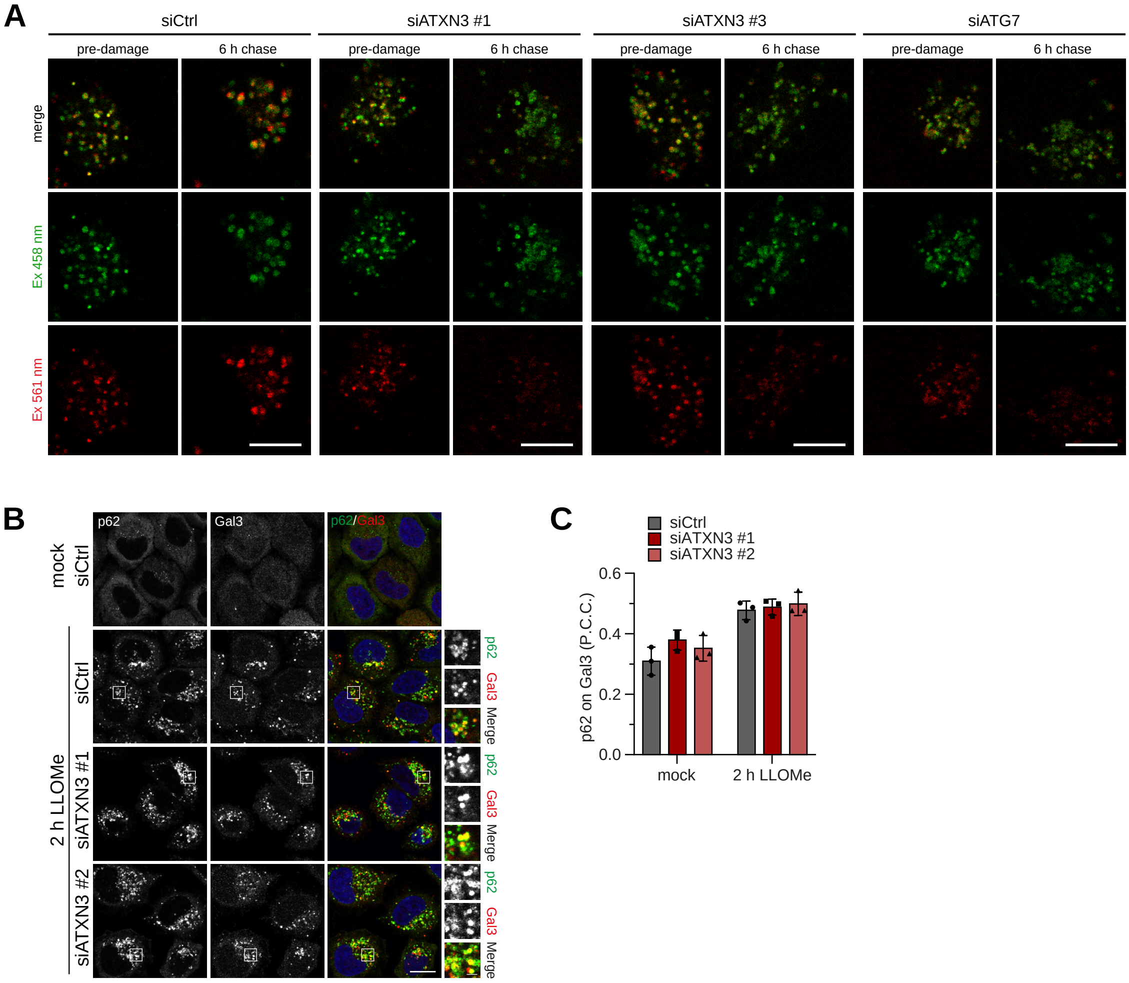


Supplementary Figure 3

**Supplementary Figure 3 (related to Fig. 4) ATXN3 is required for completion of lysophagy.**

(**A**) Micrographs corresponding to TMEM192-mKeima lysophagy assay in Fig. 4A as indicated. Scale bar, 10 µm.

(**B**) ATXN3 depletion does not affect recruitment of SQSTM1/p62 to damaged lysosomes. HeLa cells treated with indicated siRNAs were mock or LLOMe-treated and fixed at 2 h after washout. Cells were immuno-stained for Gal3 and p62. Scale bar, 15 µm.

(**C**) Quantification of (B), n=3 biological replicates with >70 cells per condition per experiment. Co-localization of p62 and Gal3 was automatically quantified with P.C.C. The graph presents the mean ± SD of three independent experiments with 79 or more cells per condition. Significance was tested by Two-way ANOVA with Dunnet's multiple comparison test, and no significant differences were found.

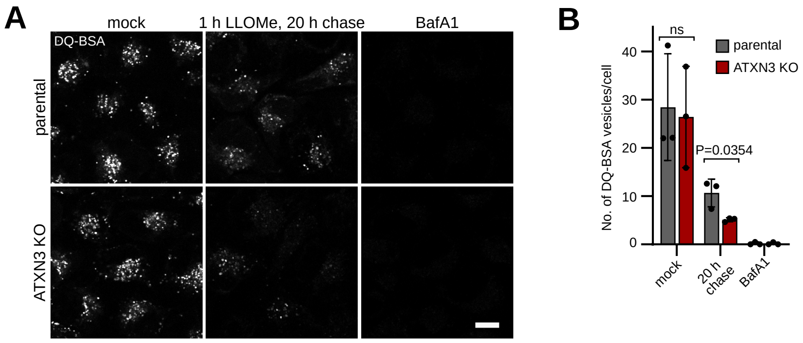


Supplementary Figure 4

**Supplementary Figure 4 (related to Fig. 5). ATXN3 is required for restoration of lysosomal degradative capacity.**

(**A**) DQ-BSA assay for proteolytic activity in lysosomes. HeLa parental and ATXN3 knockout cells were LLOMe-treated for 1 h. DQ-BSA was added 13 h after washout and DQ-BSA fluorescence in lysosomes imaged at a total of 20 h after LLOMe washout. Scale bar, 10 µm. (**B**) Quantification of (A), n=3 biological replicates with >40 cells per condition per experiment. Mixed effects analysis with Tukey's multiple comparison test. Error bars represent mean with S.D.

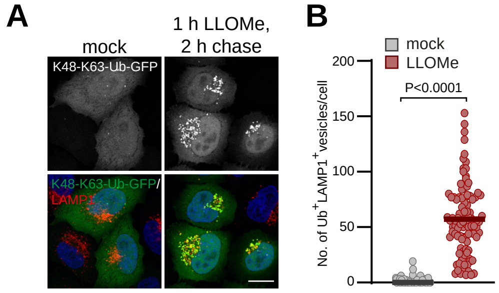


Supplementary Figure 5

**Supplementary Figure 5 (related to Fig. 6). Damaged lysosomes are modified with K48-K63-branched ubiquitin chains.**

(**A**) HeLa cells expressing NbSL3.3Q-GFP specific for K48-K63-branched chains were mock or LLOMe-treated, fixed and immuno-stained for LAMP1. Note prominent localization of NbSL3.3Q-GFP on lysosomes specifically after induction of lysosome damage. Scale bar, 15 μm.

(**B**) Quantification of (A), n=3 biological replicates with >20 cells per condition per experiment. Statistical significance was calculated with unpaired t-test. Lines represent mean.
